## Supplemental figures S1-S16 for "Ribonucleotide Reductase Subunit Switching in Hepatoblastoma Drug Response and Relapse"

Suppl Figure S1

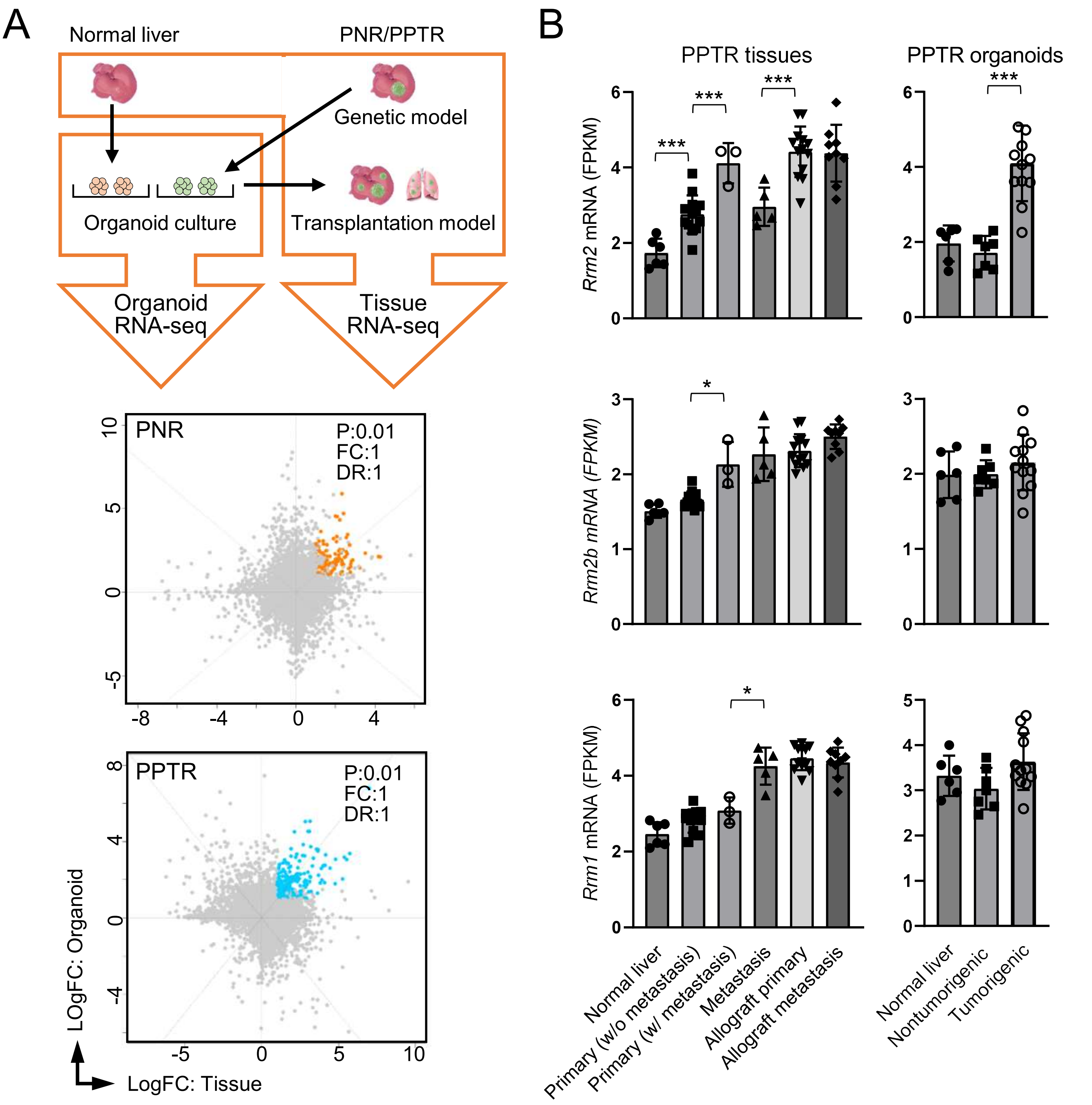


**Supplemental Figure S1. RRM2 is associated with the disease progression in PPTR mouse model of HCC.**

(A) Flow chart for the comparative transcriptomic analysis of the tumors and organoids from the PNR and PPTR mouse models.

(B) Quantitative comparison of the expression of three RNR subunits in PPTR tumor tissues (N = 6, 13, 3, 5, 13, 9, respectively, for the six groups presented) and organoids (N = 6, 7, 12, respective, for the three groups presented) in the indicated groups.

Suppl Figure S2

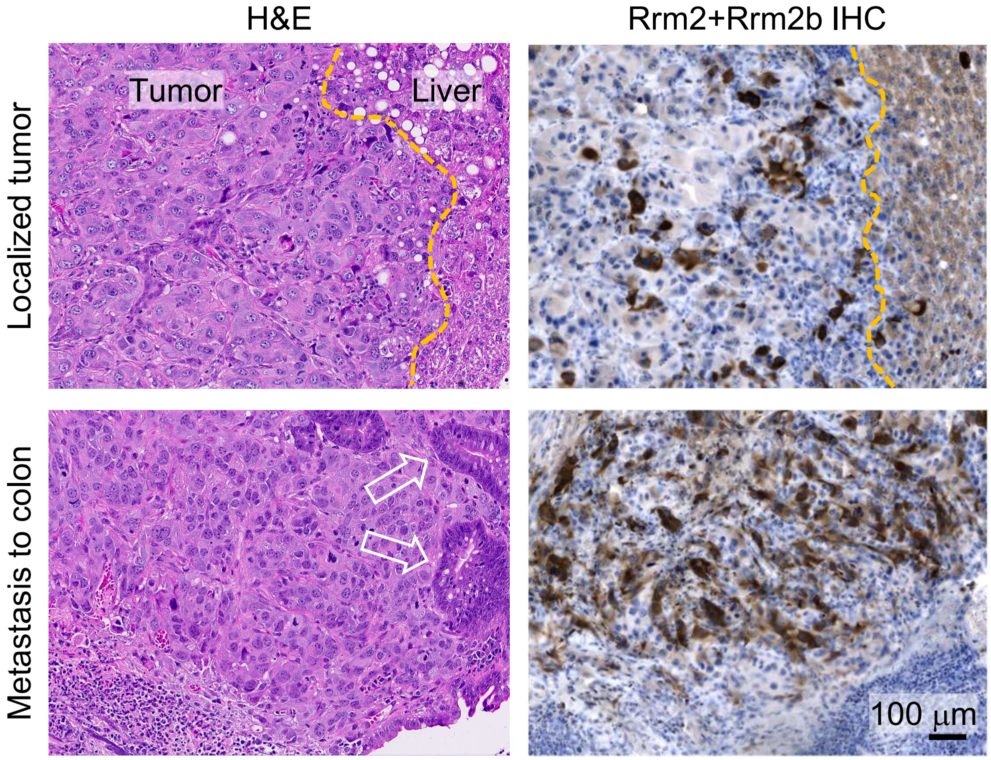


**Supplemental Figure S2. Combined RRM2 and RRM2B levels are elevated in metastatic PNR tumors compared to localized tumors.**

H&E (left) and RRM2+RRM2B IHC staining (right) on serial sections of a localized and a metastatic PNR tumor. Dotted lines: tumor border; arrows: colonic polyps embedded in the tumor. All images share the same 100 µm scale bar.

Suppl Figure S3

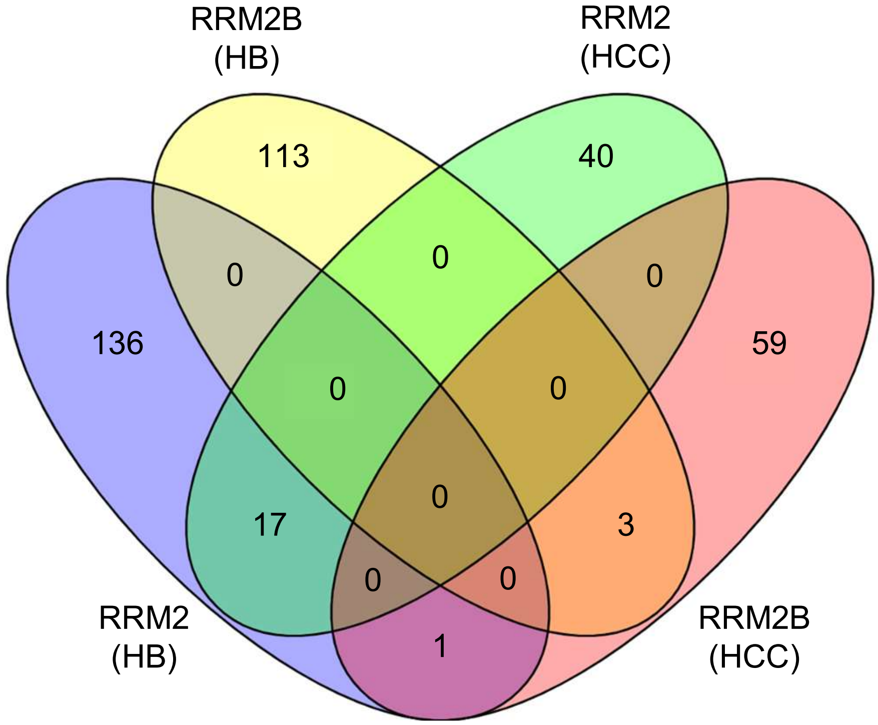


**Supplemental Figure S3. Limited overlap of the *RRM2* and *RRM2B* hub genes between HB and HCC patient tumors.**

Venn plot showing the overlap of the four indicated *RRM2* and *RRM2B* hub gene lists identified in HB and HCC patient tumors.

Suppl Figure S4

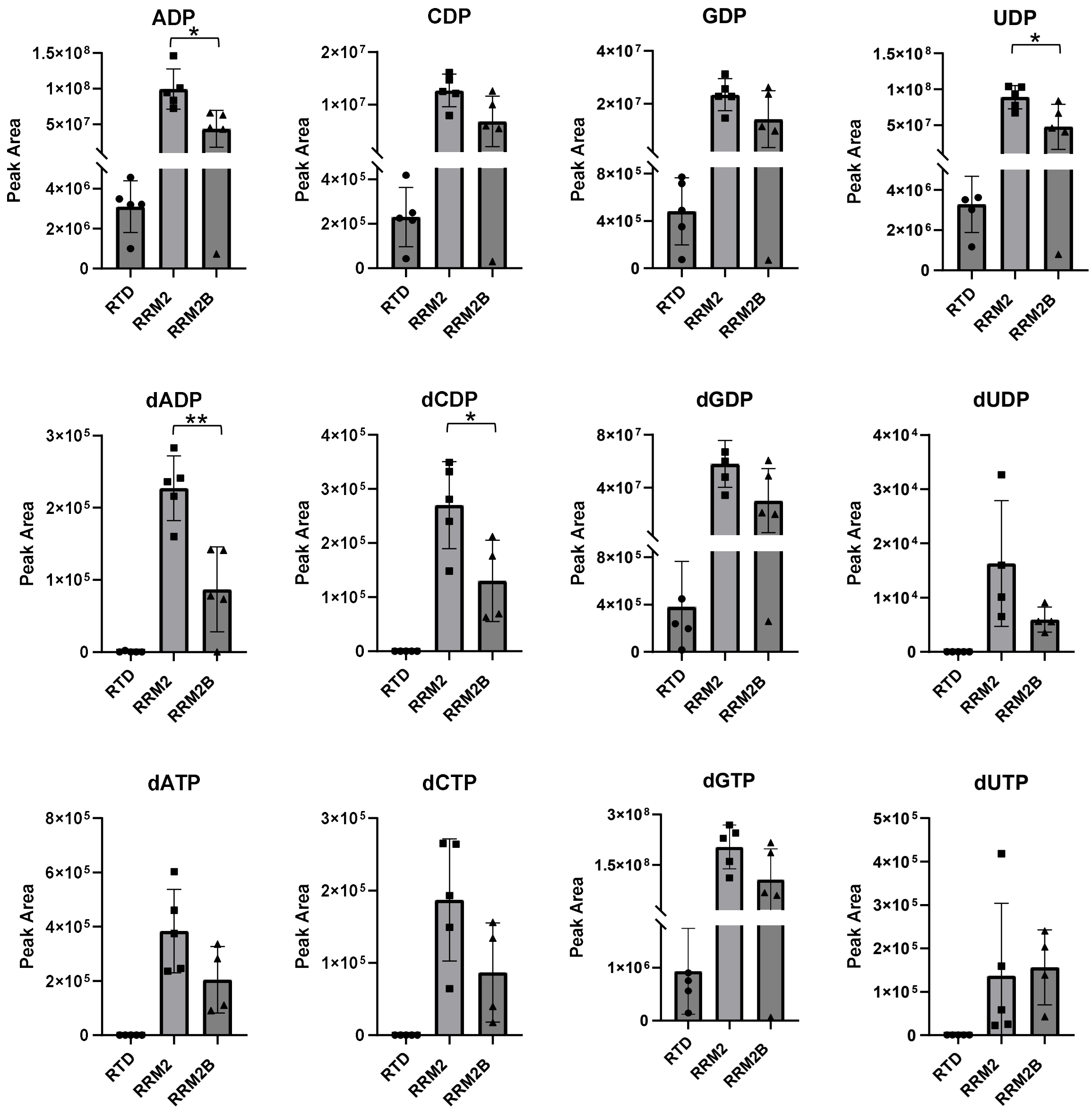


**Supplemental Figure S4. RRM2 has higher RNR enzymatic activity than RRM2B in HepG2 cells.**

Quantitative analysis of nucleotide levels in *tdT*, *RRM2$^{OE}$*, and *RRM2B$^{OE}$* HepG2 cells using targeted liquid chromatography/mass spectrometry (biological replicates: n=5 per group).

### Suppl Figure S5

A

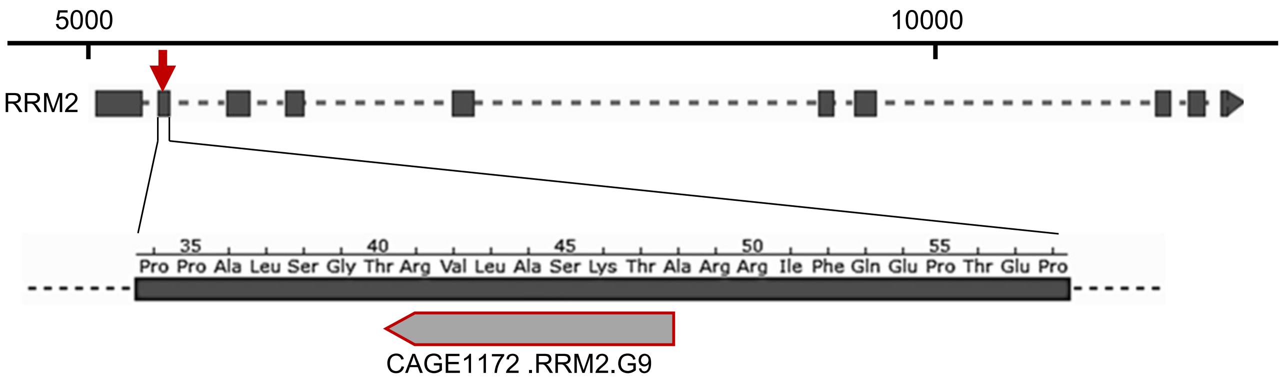


B

| Indel Ranking | bp change | Clone% |
|---|---|---|
| **#1** | **0** | **13.9%** |
| #2 | -1 | 9.2% |
| **#3** | **-18** | **7.2%** |
| **#4** | **-6** | **5.2%** |
| **#5** | **-12** | **4.5%** |
| **#6** | **-21** | **4.5%** |
| **#7** | **-24** | **4.2%** |
| #8 | -13 | 3.7% |
| **#9** | **-3** | **3.7%** |
| #10 | 1 | 3.0% |
| | Total indel | 86.10% |
| | **Total in-frame indel** | **43.2%** |

C

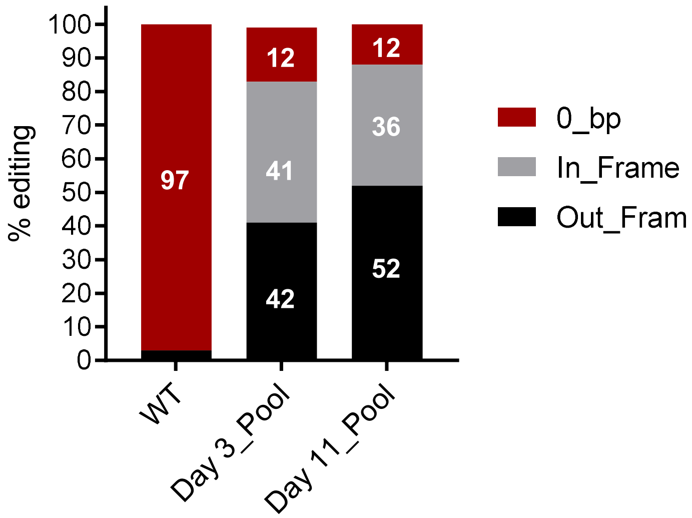


**Supplemental Figure S5. RRM2 is an essential gene in HepG2 cells.**

(A) The position and sequence of *RRM2* guide RNA.

(B) Indels after attempted knockout of *RRM2* in <u>*RRM2B*</u>$^{OE}$ HepG2 cells.

(C) Quantitative analysis of *RRM2* indels composition on Day 3 and Day 11 of in <u>*RRM2B*</u>$^{OE}$ HepG2 cells.

Suppl Figure S6

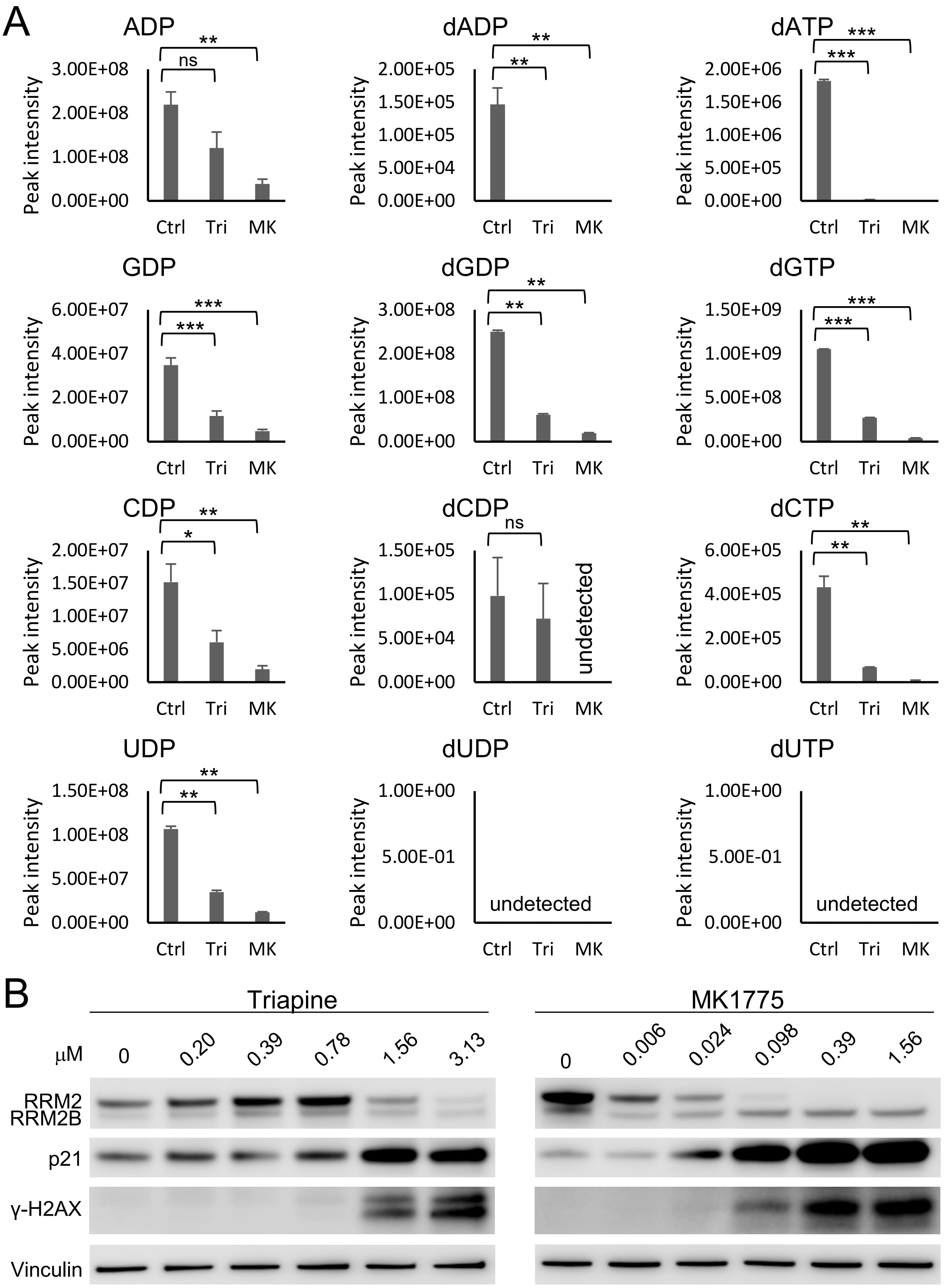


**Supplemental Figure S6. RRM2 inhibition in HepG2 cells leads to nucleotide reduction, cell cycle arrest and DNA damage response.**

(A) Quantitative analysis of nucleotide levels in HepG2 cells treated with control (ctrl, DMSO), triapine (Tri, 3.125 μM), and MK1775 (MK, 0.39 μM) via targeted liquid chromatography/mass spectrometry (biological replicates: n=5 per group).

(B) Immunoblotting of the indicated proteins in HepG2 cells treated with triapine and MK1775 at the indicated concentration.

Suppl Figure S7

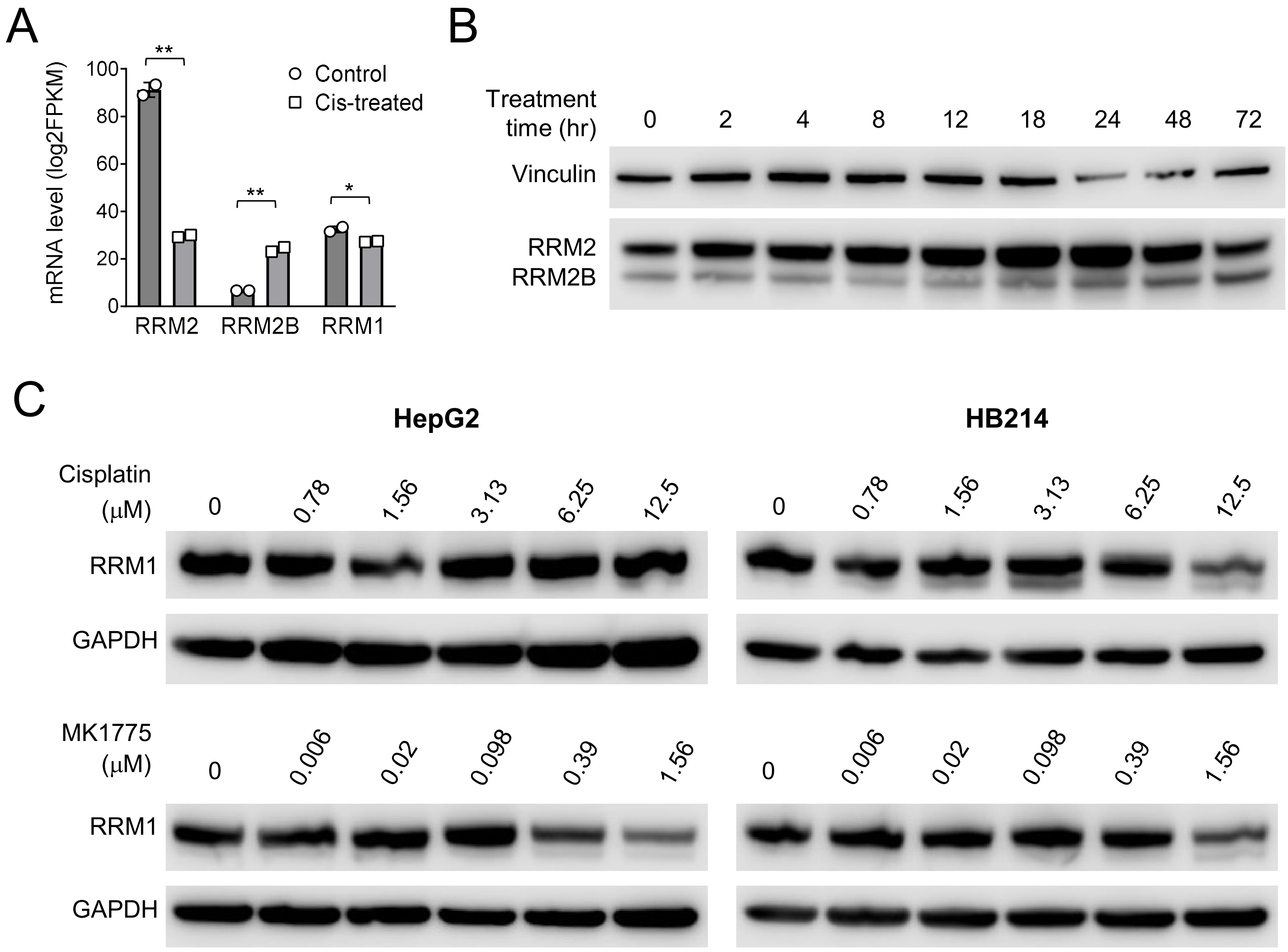


**Supplemental Figure S6. Drug treatment induces different gene expression changes to the three RNR subunits in HepG2 cells.**

(A) RNR subunits mRNA levels detected by RNAseq in HepG2 cells treated with cisplatin (biological replicates: n=2 per group).

(B) A time-course study of RRM2 and RRM2B protein levels in cis-treated HepG2 cells by immunoblotting.

(C) RRM1 immunoblotting in HepG2 and HB214 cells treated with cisplatin and MK1775.

### Suppl Figure S8

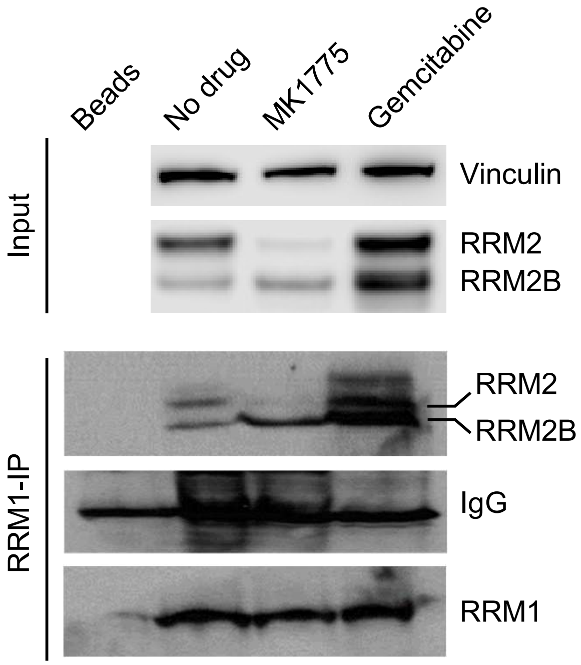


**Supplemental Figure S7. RRM2B-RRM1 complex is the dominant RNR complex in drug-treated HepG2 cells.**

RRM1 co-IP assay using MK1775- and gemcitabine-treated HepG2 cells.

### Suppl Figure S9

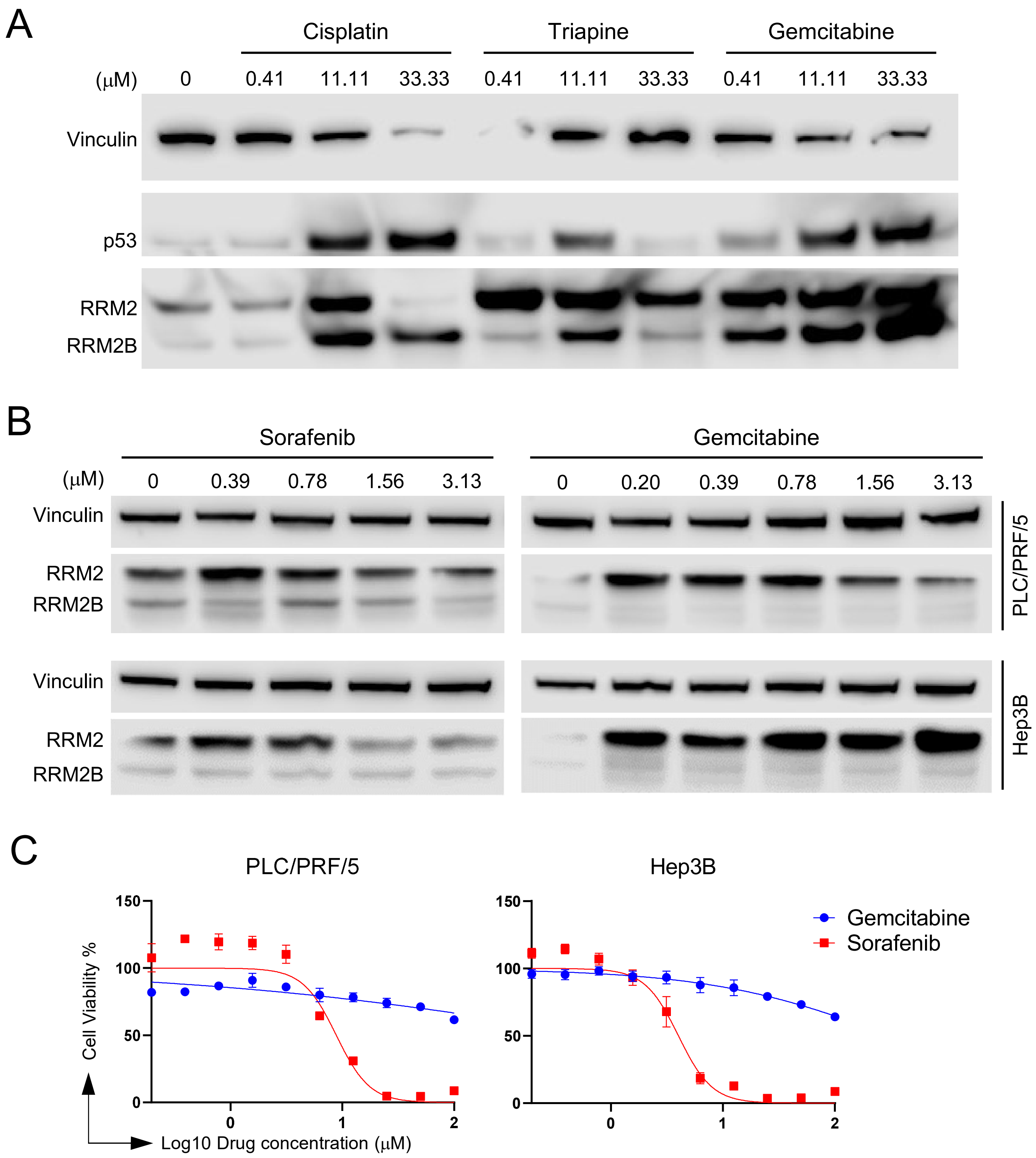


**Supplemental Figure S8. Drug-induced <u>RRM2B</u> upregulation in HB cells coincides with p53 induction.**

(A) Immunoblot of p53, RRM2 and <u>RRM2B</u> in HepG2 cells treated with the indicated drugs.

(B) RRM2 and <u>RRM2B</u> immunoblotting in two *TP53*-mutant HCC cell lines, PLC/PRF/5 and Hep3B, treated with sorafenib and gemcitabine.

(C) Dose response curves of PLC/PRF/5 and Hep3B cells treated with sorafenib and gemcitabine. All drug curves represent three technical replicates. All assays were biologically repeated for three times.

### Suppl Figure S10

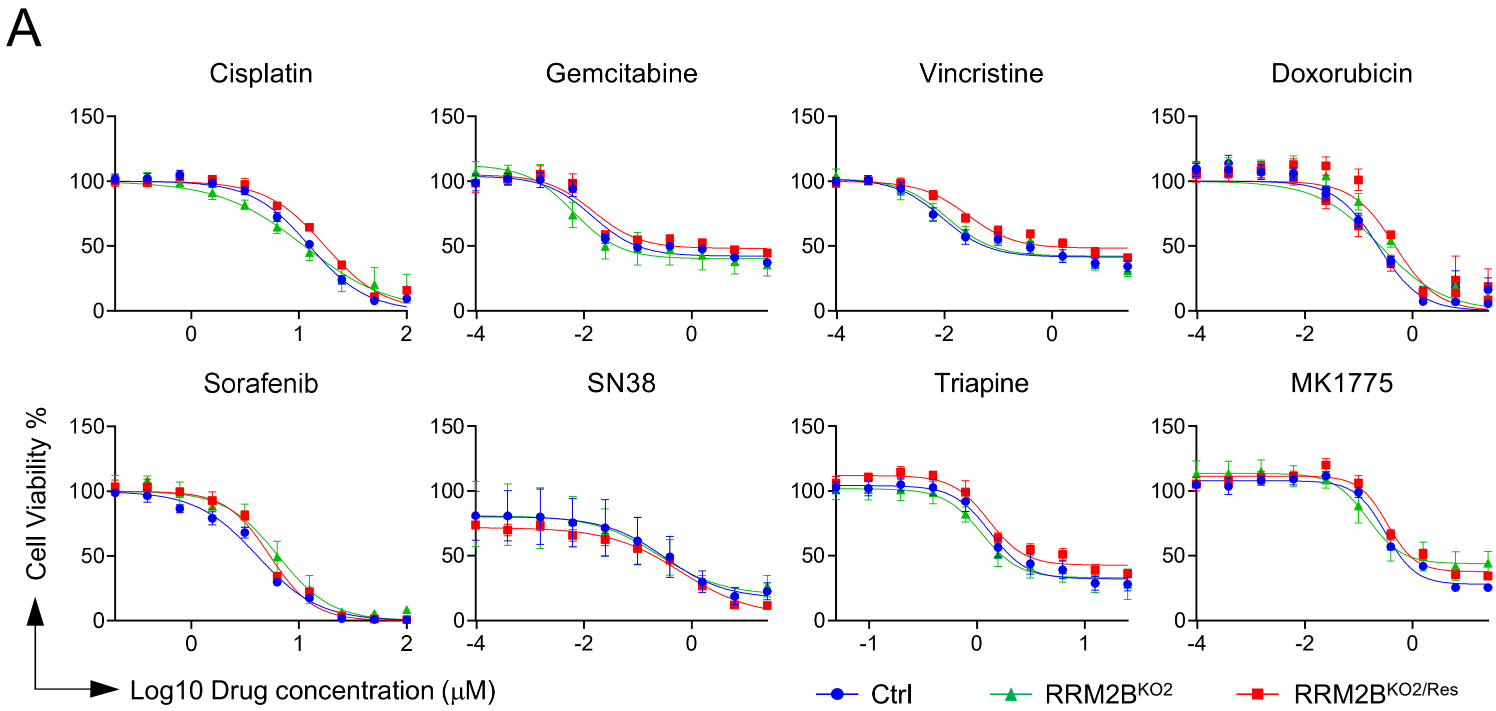


B

| | Cisplatin | Gemcitabine | Vincristine | Doxorubicin | Sorafenib | SN38 | Triapine | MK1775 |
|---|---|---|---|---|---|---|---|---|
| $IC_{50}$: Ctrl | 11.27 | 1.354 | 1.305 | 0.2427 | 4.62 | 0.2805 | 3.906 | 1.801 |
| $IC_{50}$: RRM2B$^{KO2}$ | 11.21 | 0.4569 | 0.5218 | 0.2508 | 6.416 | 0.2676 | 3.501 | 3.51 |
| $IC_{50}$: RRM2B$^{KO2/Res}$ | 17.5 | 2.333 | 2.008 | 0.4716 | 5.383 | 0.1512 | 7.082 | 2.939 |
| *P* value: RRM2B$^{KO2}$ vs. ctrl | 0.934 | 0.0164 | 0.0737 | 0.9283 | <.0001 | 0.9174 | 0.4085 | 0.0538 |
| *P* value: RRM2B$^{KO2/Res}$ vs. RRM2B$^{KO2}$ | <.0001 | 0.0005 | 0.0097 | 0.0007 | 0.0006 | 0.2176 | <.0001 | 0.5846 |

**Supplemental Figure S9. Drug response curves of control, *RRM2B*$^{KO2}$ and *RRM2B*$^{KO2/Res}$ HepG2 cells.**

(A) The dose-response curves of control (wildtype), *RRM2B*$^{KO2}$ and *RRM2B* $^{KO2/Res}$ HepG2 cells to the indicated drugs. All drug curves represent three technical replicates. All assays were biologically repeated for three times.

(B) List of the drug $IC_{50}$ values and their comparisons between control (wildtype), *RRM2B*$^{KO2}$ and *RRM2B* $^{KO2/Res}$ HepG2 cells. Extra Sum of Square F test. A *P* value < 0.05 is considered statistically significant.

Suppl Figure S11

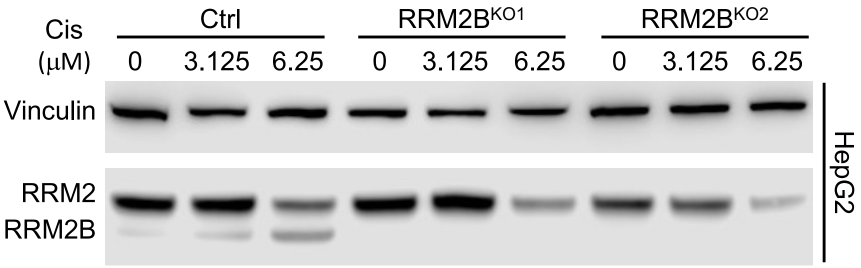


**Supplemental Figure S10. Confirmation of the lack of *RRM2B* induction in *RRM2B$^{KO}$* HepG2 cells.**

RRM2 and RRM2B immunoblotting in untreated and cisplatin-treated *RRM2B$^{KO}$* HepG2 cells.

### Suppl Figure S12

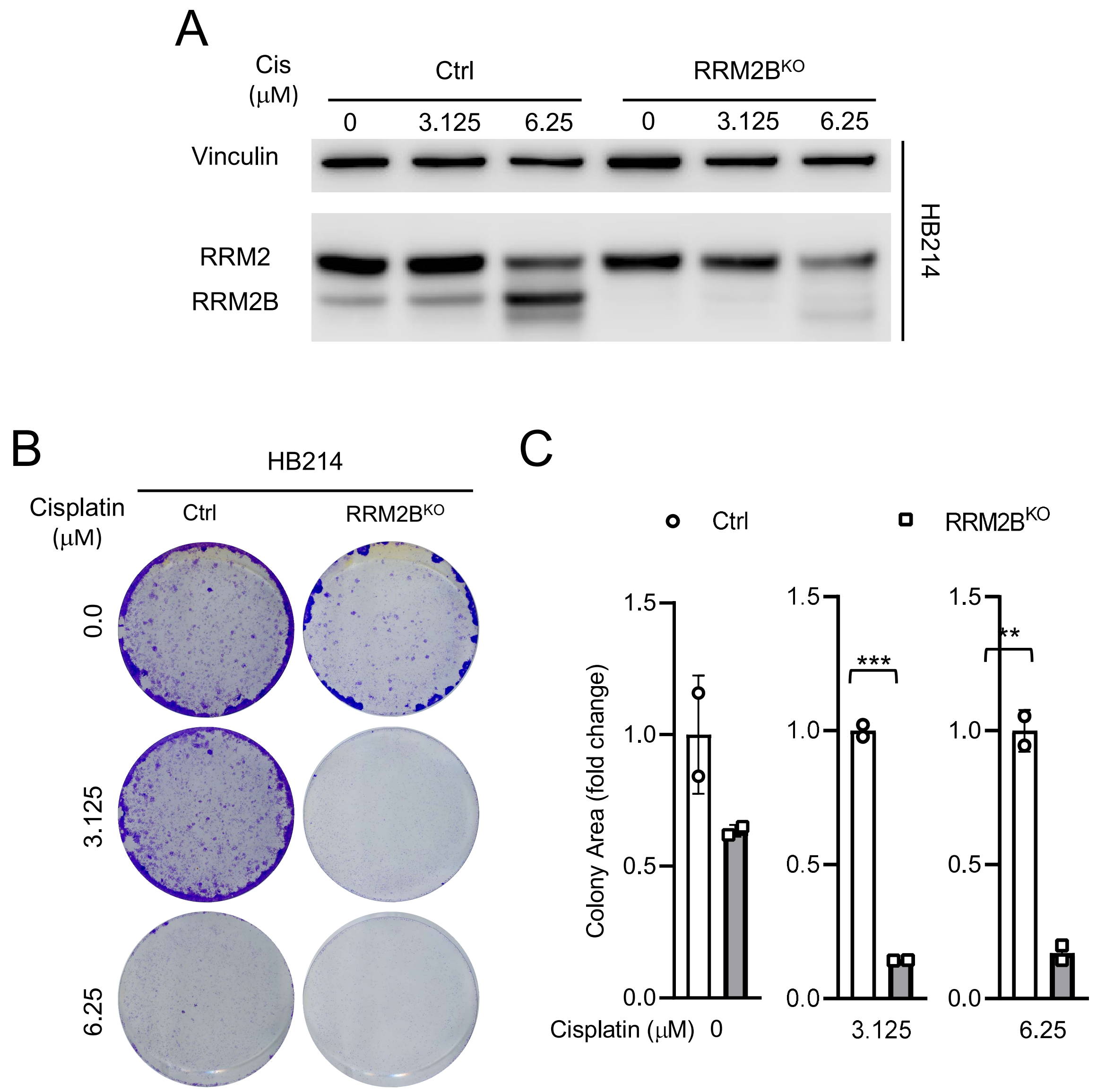


**Supplemental Figure S11. RRM2B supports post-drug treatment recovery of HB214 cells in vitro.**

(A) RRM2 and RRM2B immunoblotting in untreated and cis-treated *RRM2B*$^{KO}$ HB214 cells.

(B) 12-day colony formation assay of the control and *RRM2B*$^{KO}$ HB214 cells post the indicated treatment.

(C) Quantitative analysis of area occupied by cells in **(B)** (biological replicates, n = 2 per group).

### Suppl Figure S13

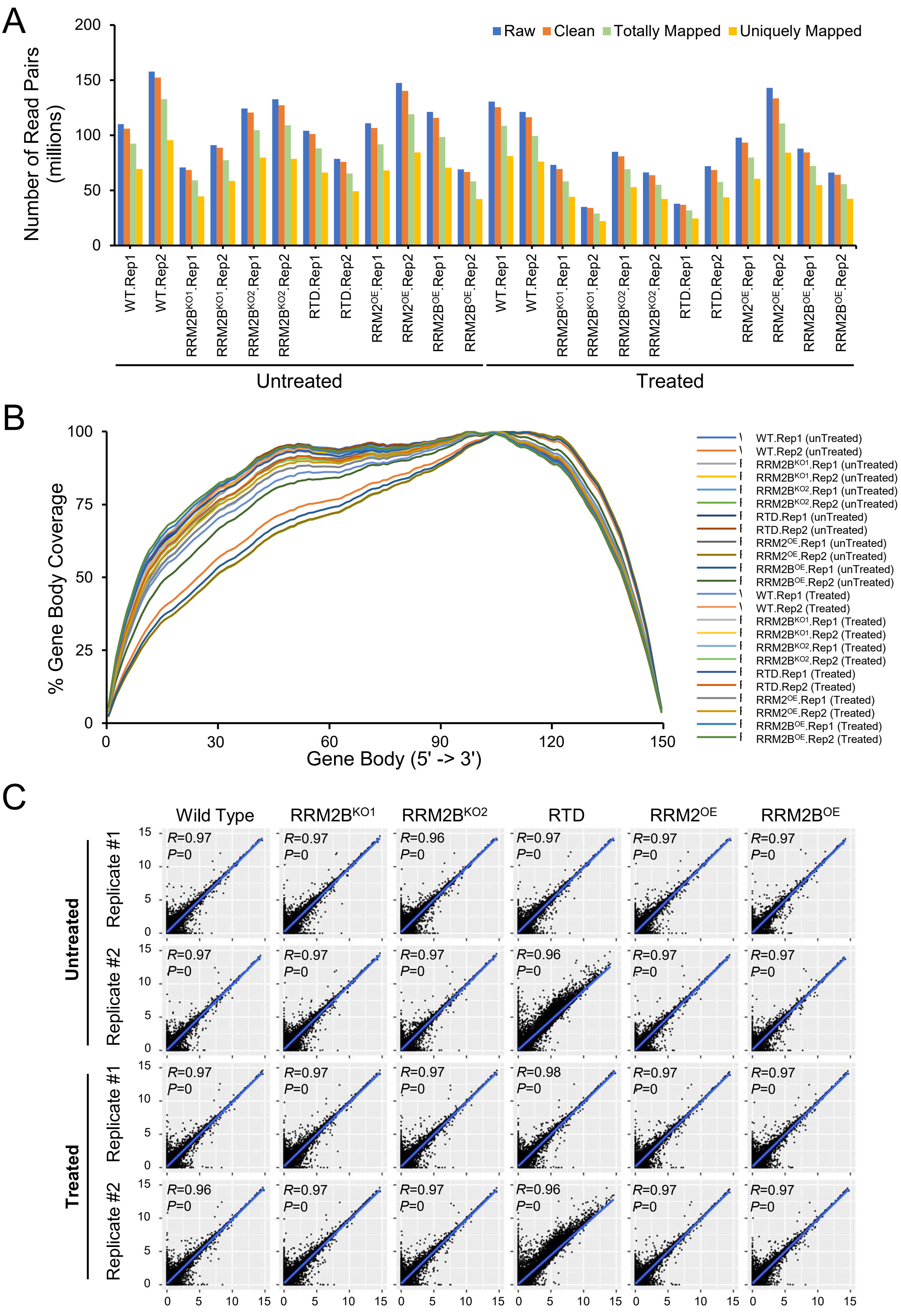

**Supplemental Figure S13. Quality assessment of HepG2 cell transcriptomic analyses.**

(A) Bar plot showing the alignment statistics of the RNA-Seq data.
(B) The gene body coverage statistics at the resolution of 150 bins per transcript.
(C) The accuracy evaluation of gene expression quantification by RSEM. The Spearman correlation coefficient and P-value were calculated from the genes co-identified by RSEM and Salmon using the stats R package (v3.6.1).

Suppl Figure S14

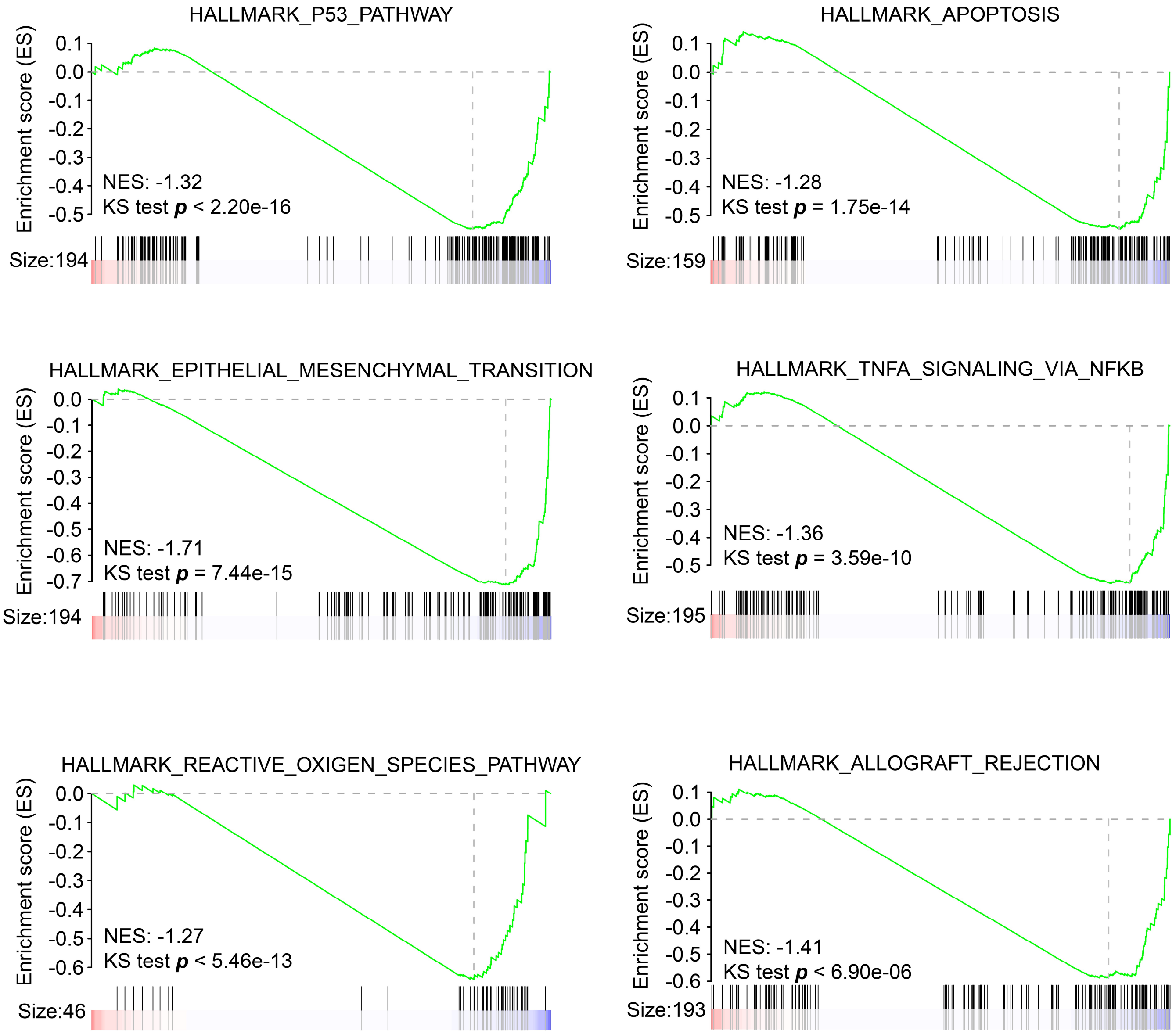


**Supplemental Figure S13. GSEA plots of six main RRM2B-involved pathways.**

Suppl Figure S15

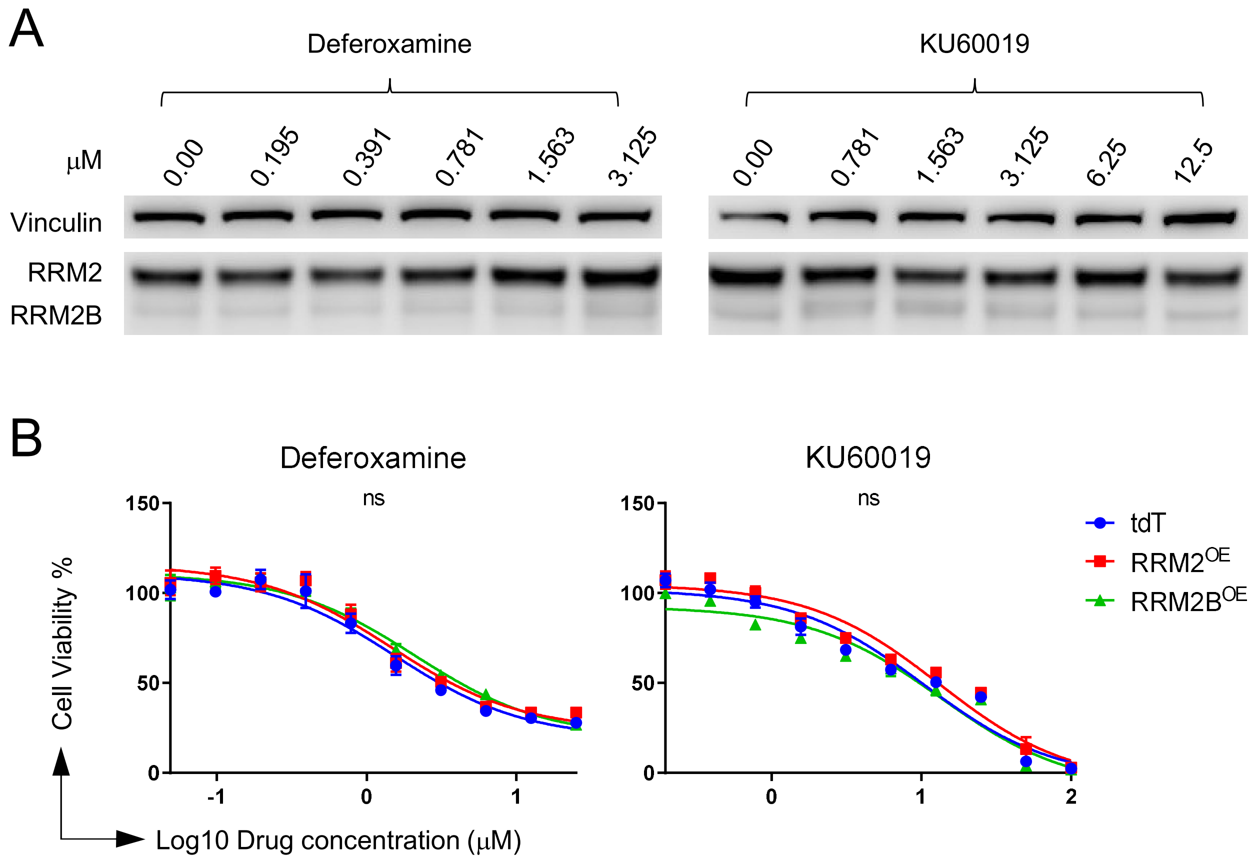


**Supplemental Figure S14. Two putative RRM2B inhibitors deferoxamine and KU60019 have no RRM2B-specific inhibition in HepG2 cells.**

(A) Immunoblot for RRM2 and RRM2B in HepG2 cells treated with deferoxamine and KU60019.

(B) Dose response curves of *tdT*, *RRM2*$^{OE}$, and *RRM2B*$^{OE}$ HepG2 cells to deferoxamine and KU60019. Extra Sum of Square F test; ns: not significant (*P* value > 0.05). All drug curves represent three technical replicates. All assays were biologically repeated for three times.

### Suppl Figure S16

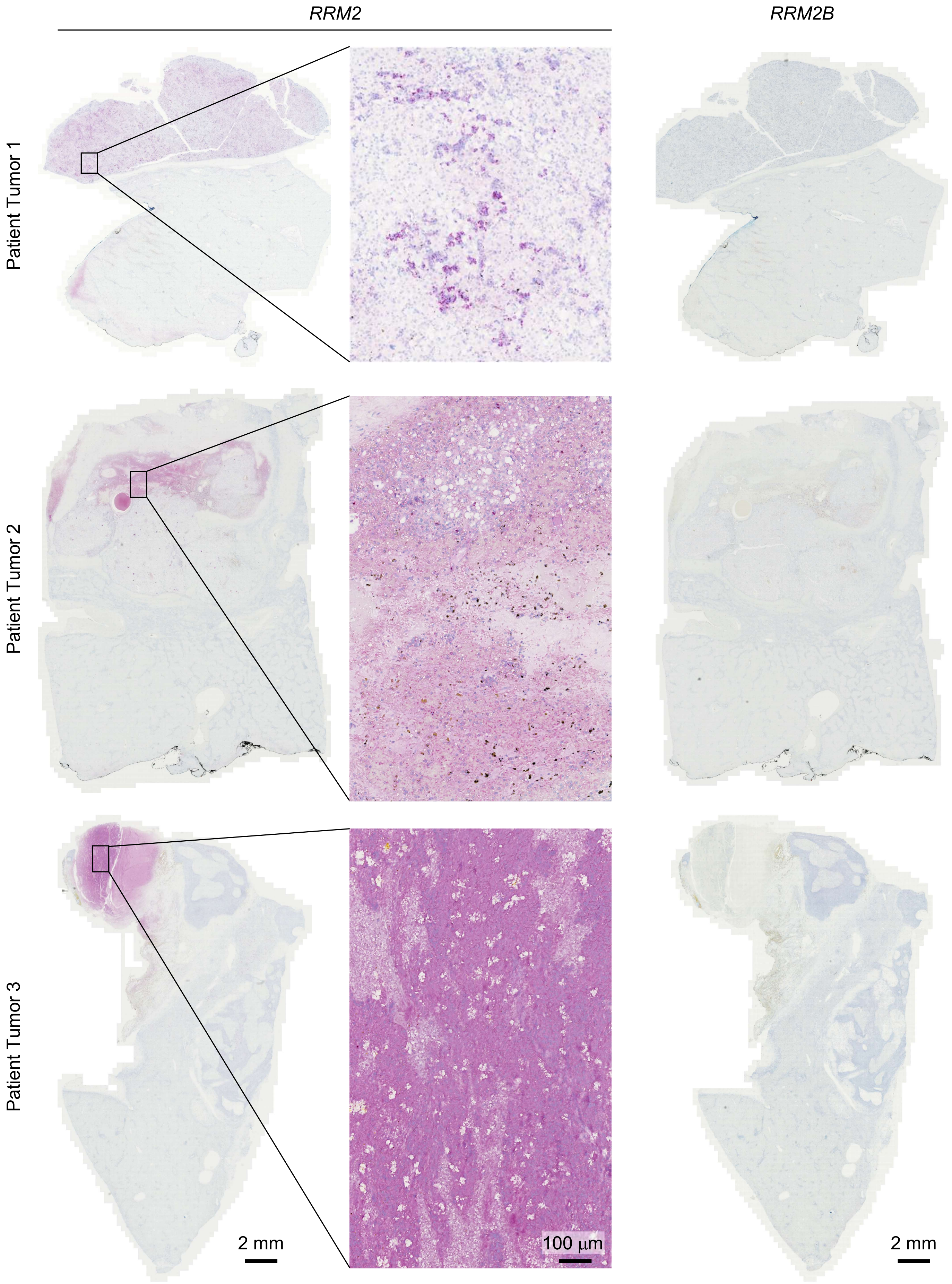


**Supplemental Figure S15. *RRM2* and *<u>RRM2B</u>* RNAscope staining on three primary HB patient tumors.** Images on the same column share the same scale bar.
